# Developmental Reorganization of Whole-Body Muscle Synergies During Overarm Throwing in Children

**DOI:** 10.1101/2025.10.08.681079

**Authors:** Takuya Murakami, Ayane Kusafuka, Taishi Okegawa, Daiki Yamasaki, Yuto Sakakibara, Nadaka Hakariya, Naotsugu Kaneko, Hiroki Saito, Atsushi Sasaki, Kimitaka Nakazawa

## Abstract

Overarm throwing is a uniquely human skill that requires precise whole-body coordination. Although throwing behavior emerges early in childhood, the neuromuscular mechanisms that support its development remain poorly characterized. Here, we provide novel evidence for the developmental reorganization of whole-body muscle synergies during maximum-effort throwing in preschool-aged (PS) and school-aged (SA) children. Electromyography was recorded from 16 muscles, and non-negative matrix factorization was applied to extract low-dimensional coordination modules (muscle synergies). We compared ball speed, number of synergies, synergy structure, and temporal consistency between groups. Ball speed was significantly higher in SA than PS (33.6 ± 10.2 vs. 21.4 ± 6.2 km/h, p < 0.05), reflecting improved performance. Yet, the number of synergies did not differ (PS: 6.0 ± 1.1; SA: 6.4 ± 1.3, p > 0.05), suggesting that the dimensionality of coordination is largely established by the preschool years. Instead, developmental improvements were driven by structural and temporal reorganization: trunk- and upper-limb synergies merged into a single module in SA, reflecting improved postural integration, while a bilateral soleus-dominant synergy fractionated into lateralized modules, reflecting increased lower-limb specialization. Moreover, the temporal variability of synergy activation was reduced in SA (p < 0.01), indicating that movement sequences became more precise and stable with development. These findings reveal that early gains in throwing arise not from expanding synergy number but from reorganizing their structure and sharpening temporal coordination, offering mechanistic insight into how complex whole-body skills are refined during childhood.

**Significance Statement:** Throwing is a hallmark of human motor behavior, requiring precise sequencing of whole-body muscle activity. Yet how children develop this ability has remained unclear. By applying muscle synergy analysis to electromyographic recordings of preschool and school-aged children performing maximum-effort throws, we found that improvements in performance were not due to an increase in synergy number but rather to structural reorganization and greater temporal precision. Specifically, trunk and upper-limb modules merged, lower-limb modules fractionated, and activation timing became more consistent. These results identify merging and fractionation as complementary mechanisms supporting developmental refinement of motor skills. More broadly, they provide a mechanistic framework for understanding how complex whole-body actions are acquired and offer markers for pediatric training and rehabilitation strategies.

## Introduction

The ability to throw objects with speed and accuracy is a uniquely human skill that depends on sophisticated whole-body muscle coordination. Throwing has played a pivotal role throughout human history, from hunting in ancestral foraging societies to serving as a central skill in modern sports such as baseball and cricket (1). Interestingly, throwing behavior emerges spontaneously in early childhood and develops even without formal instruction (2), suggesting deeply embedded neural foundations with evolutionary significance in the human motor system. Overarm throwing, therefore, serves as an ideal model for investigating how humans develop and refine whole-body coordination. Understanding its developmental trajectory may provide insight into the maturation of neuromotor control.

To execute such complex movements, the central nervous system (CNS) must generate precisely coordinated motor commands across hundreds of muscles and thousands of motor units. A prevailing hypothesis is that the CNS simplifies this high-dimensional control problem through a modular strategy, commonly described as muscle synergies: coordinated patterns of muscle activation that reduce dimensionality of motor control (3). These muscle synergies can be extracted from electromyographic (EMG) data using non-negative matrix factorization (NMF), which reveals the underlying muscle coordination structures involved in task execution (4). Developmental studies of locomotion demonstrated that muscle synergies follow a staged trajectory: newborns express two basic synergies, toddlers acquire two additional ones, and these four modules are refined but largely conserved through adulthood (5). Running similarly shows an increase in modular complexity, from approximately six synergies in preschoolers to seven in sedentary adults (6).

While developmental changes in locomotor muscle synergies are well characterized, much less is known about how coordination emerges and is refined across development during complex upper-limb tasks such as throwing. A recent study focusing only on upper-limb muscles reported that the number of throwing-related synergies remained stable from preschool to school age, but their structure reorganized across development (7). However, throwing fundamentally requires precise integration of upper-limb, trunk, and lower-limb actions to generate power and transfer energy through the kinetic chain (8). Importantly, the transition from preschool (typically under 6–7 years of age) to the early school years represents a pivotal developmental window when coordination and fundamental motor skills become more organized (9). Indeed, classic observational work showed that the fundamental throwing pattern emerges by around six and a half years of age, with further refinements in speed and coordination continuing through approximately twelve years (2). Thus, examining whole-body muscle synergies during this stage is critical to uncovering how children reorganize neuromuscular modules to achieve efficient throwing. By extending muscle synergy analysis beyond the upper limb, the present study addresses a key gap and provides evidence for how whole-body coordination, including the integration of upper-limb, trunk, and lower-limb modules, is refined during development. This whole-body perspective provides a powerful framework for quantifying developmental changes, as it captures the low-dimensional modular organization underlying high-dimensional, whole-body movements.

Accordingly, the present study aimed to investigate developmental changes in whole-body muscle synergies during overarm throwing by comparing preschool-aged and elementary school-aged children. Based on locomotor studies showing both early establishment of basic modules (5) and later increases in module number during adolescence (6), as well as prior evidence of structural reorganization in throwing synergies (7), we hypothesized that (i) the number and structure of muscle synergies would differ between the two age groups, reflecting developmental refinement to enhance throwing performance (i.e., ball speed), and (ii) synergy activation timing would become more consistent across trials, indicating increased coordination efficiency as children’s movements align more precisely with key mechanical events (e.g., foot contact, trunk rotation, and ball release) during throwing (8, 10). By clarifying the developmental plasticity of throwing-related motor modules, this study seeks to advance understanding of the neural mechanisms that underlie developmental motor skill acquisition and provide a basis for evidence-based training and intervention strategies in pediatric populations.

## Results

### Throwing performance and the number of muscle synergies

We recorded maximum-effort overarm throws in 11 preschoolers (PS; mean age 62.2 ± 8.0 months) and 9 school-aged children (SA; mean age 100.3 ± 14.4 months). None of the participants had prior experience in organized sports involving overarm throwing (e.g., baseball or softball). Participants threw a ball toward a net 2 m away while surface EMG was simultaneously recorded from 16 muscles spanning the upper limbs, trunk, and lower limbs.

Ball speed, derived from high-speed camera tracking, was significantly higher in SA (33.6 ± 10.2 km/h) than in PS (21.4 ± 6.2 km/h) (p < 0.05, Figure 1A), reflecting age-related improvements in throwing performance.

**Figure 1:**
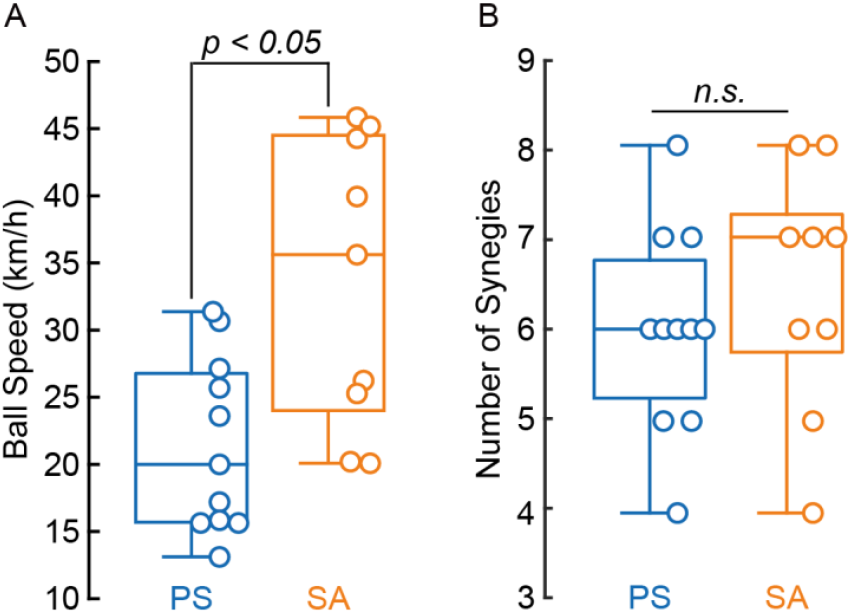
Developmental changes in throwing performance and number of synergies. (A) Ball speed during maximum-effort overarm throws in preschool (PS) and school-aged (SA) children. Each dot represents an individual participant; boxplots show the median (horizontal line), interquartile range (box), and full data range (whiskers). Ball speed was significantly higher in the SA group compared to the PS group (p < 0.05), indicating developmental improvements in throwing performance. (B) Number of muscle synergies extracted from PS and SA children. Each dot represents an individual participant. The number of synergies did not differ significantly between groups (n.s.).

To evaluate underlying coordination, EMG signals were processed (band-pass filtered, rectified, low-pass filtered, normalized, and time-interpolated) and decomposed using non-negative matrix factorization (NMF) to extract low-dimensional muscle synergies; only solutions achieving ≥ 90% variance accounted for (VAF) were retained. The average number of muscle synergies did not differ significantly between PS (6.0 ± 1.1) and SA (6.4 ± 1.3) (p > 0.05) (Figure 1B). This indicates that the overall complexity of motor coordination, as reflected by the number of synergies, remains stable between the two groups, despite differences in throwing performance.

### Developmental reorganization of synergy structure

Clustering analysis was applied to characterize differences in synergy structure between the two groups. This revealed nine muscle synergies in PS and eight in SA. Among them, six were common synergies, while three were specific to PS and two were specific to SA (Figure 2A). Synergies were classified based on whether pairwise cosine similarity exceeded 0.9.

**Figure 2:**
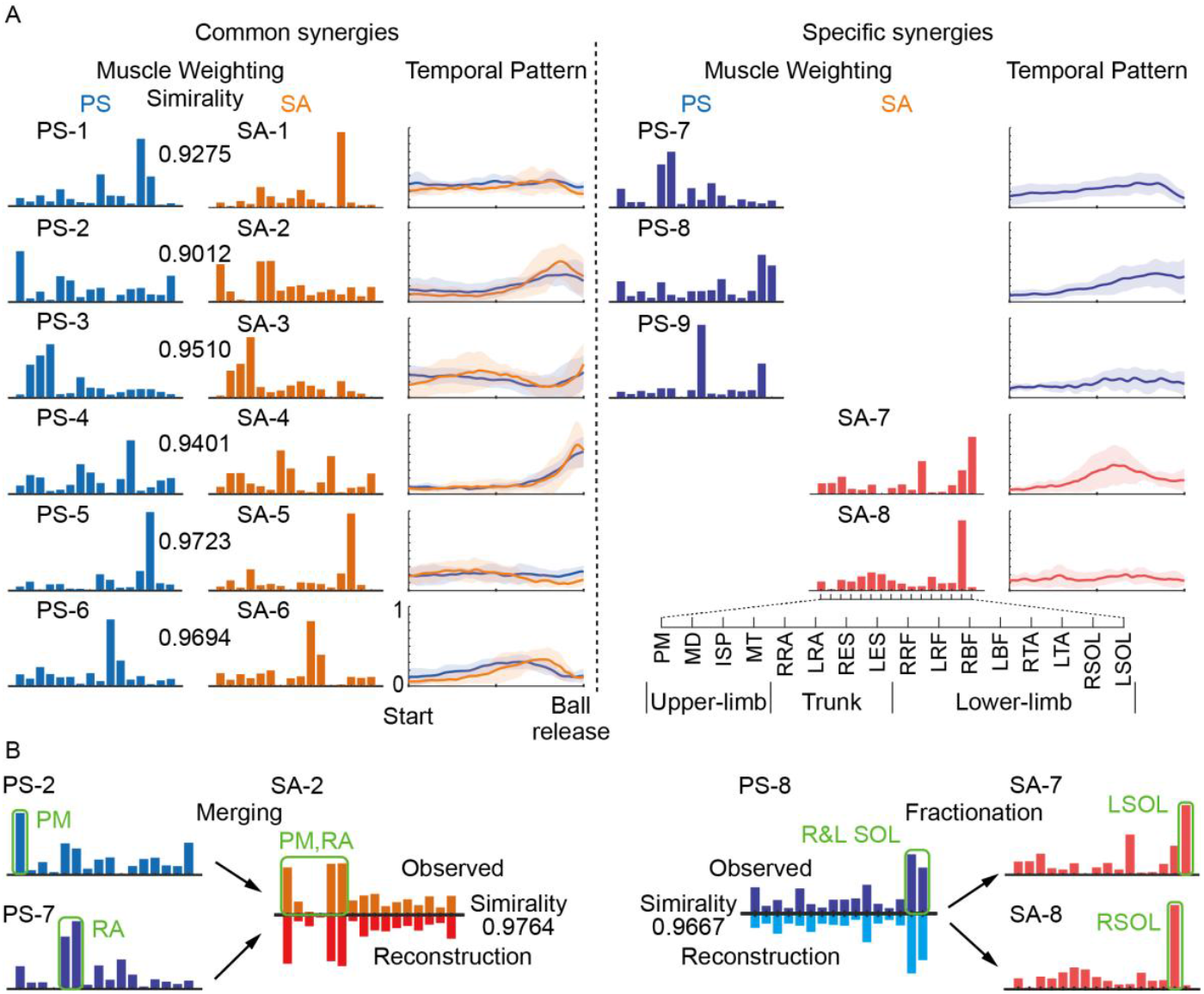
Developmental reorganization of whole-body muscle synergies during throwing. (A) Representative muscle synergy extracted from preschool (PS) and school-aged (SA) children. Six common synergies (PS-1 to PS-6; SA-1 to SA-6) were shared across both groups, while additional group-specific synergies emerged (PS-7 to PS-9; SA-7 and SA-8). Bars represent the muscle weighting categorized by anatomical region (upper limb, trunk, lower limb). Lines show the mean temporal activation profiles, with shaded areas indicating ±1 standard deviation (SD) across participants. (B) Developmental reorganization revealed by cross-group reconstruction analysis. Left: Merging—two PS synergies (PS-2: pectoralis major [PM] dominant, and PS-7: rectus abdominis [RA] dominant) combined into a single SA synergy (SA-2), indicating improved trunk-upper limb integration. Right: Fractionation—a bilateral soleus (SOL) dominant synergy in PS (PS-8) split into two distinct SA synergies (SA-7: right SOL dominant, SA-8: left SOL dominant), reflecting increased lower-limb lateralization.

We further examined cross-group reorganization using cosine similarity and reconstruction analyses. Two patterns of developmental reorganization emerged (Figure 2B):

#### Merging

In SA, a new muscle synergy (SA-2) emerged that combined two PS modules, PS-2 (pectoralis major-dominant) and PS-7 (rectus abdominis-dominant), indicating integration of trunk-upper limb coordination. The reconstruction coefficients for PS-2 and PS-7 were 0.5081 and 0.5759, respectively, and a cosine similarity between the reconstructed synergy and SA2 was 0.9764.

#### Fractionation

In contrast, PS-8 (bilateral soleus-dominant) was fractionated into two synergies in SA (SA-7, right soleus-dominant; SA-8, left soleus-dominant), indicating increased lateralization of lower-limb coordination. The reconstruction coefficients were 0.6189 and 0.4728, with a cosine similarity between the reconstructed synergy and PS-8 was 0.9667.

These findings demonstrate that, although the number of muscle synergies remained unchanged, their structural organization underwent qualitative reconfiguration across development.

### Temporal consistency of synergy activation

To evaluate the timing stability of each synergy activation, we calculated the centroid of activity (CoA) for each temporal activation profile. The standard deviation of CoA across trials was significantly lower in SA (8.0% ± 2.3%) than in PS (11.2% ± 1.7%; p < 0.01, Figure 3B). This reduction in variability indicates that synergy activation timing became more consistent across trials in the school-aged group. Representative activation profiles illustrate this effect, with narrower dispersion of CoA in SA compared to PS (Fig. 3A).

**Figure 3:**
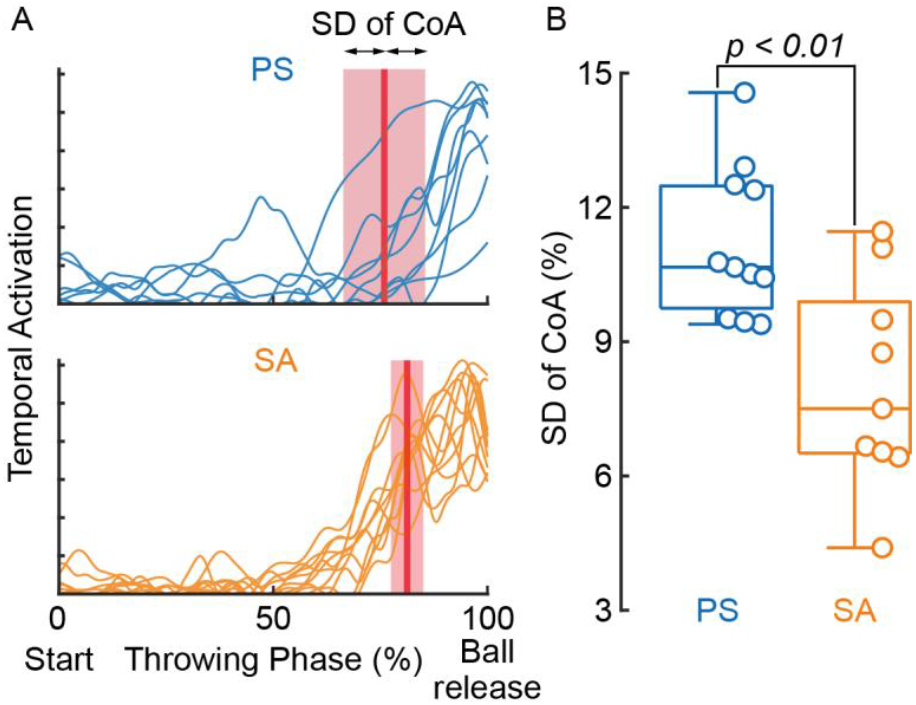
Developmental refinement of temporal consistency in synergy activation. (A) Temporal activation profiles of one representative synergy from preschool (PS, blue) and school-aged (SA, orange) participants. The vertical line indicates the centroid of activity (CoA), and the shaded area shows its standard deviation (SD) across trials. Broader dispersion in PS reflects greater variability in activation timing, whereas SA shows narrower variability around the CoA. (B): Group comparison of the SD of CoA for each participant, reflecting temporal variability of synergy activation. Each dot represents an individual participant. The SA group exhibited significantly lower variability than the PS group (p < 0.01), indicating that synergy activation timing becomes more consistent with development.

These results suggest that developmental improvements in throwing performance were supported not by increased synergy number but by structural reorganization and enhanced temporal precision.

## Discussion

In this study, we examined developmental changes in whole-body muscle coordination during overarm throwing by analyzing EMG activity from 16 upper-limb, trunk, and lower-limb muscles in PS and SA. Using NMF, we extracted low-dimensional muscle synergies and compared their number, structure, and temporal organization across groups. The main findings were: (i) throwing performance, as indexed by ball speed, progressively improved from the PS to SA; (ii) the number of muscle synergies did not differ between groups; (iii) structural reorganization was evident through merging and fractionation of modules; and (iv) synergy activation timing became more consistent in SA. Together, these results demonstrate that developmental improvements in throwing performance depend less on increasing the number of synergies and more on reorganization of their structures and refinement of temporal coordination.

### Developmental reorganization of muscle synergy structures

Contrary to our initial hypothesis, the number of muscle synergies did not differ significantly between PS and SA. This suggests that fundamental primitives of muscle coordination for throwing are already present by the preschool years. Importantly, however, the present study focused on a relatively narrow developmental window from preschool to school age. Thus, the absence of group differences in synergy number should not be taken to imply that the repertoire remains fixed throughout life. It remains possible that synergy number increases later in development, particularly during adolescence, when musculoskeletal growth, neural maturation, and structured training may further expand the motor repertoire. Indeed, studies of locomotion have reported such developmental increases: toddlers initially express four basic locomotor synergies, which are subsequently refined and expanded into the adult repertoire (5), while the number of running synergies increases from approximately six in preschool children to seven or more in sedentary adults (6). Our results, therefore, capture an early stage of refinement in which qualitative reorganization occurs without quantitative increases in synergy number.

One notable reorganization observed in the present study was the merging of upper-limb and trunk modules: the pectoralis major-dominant synergy (PS-2) and rectus abdominis-dominant synergy (PS-7) combined into a single module in school-aged children (SA-2). This integration reflects improved trunk–upper-limb coordination, a critical factor for generating greater ball speed (11, 12). The developmental shift from segregated upper-limb and trunk muscles to a co-activated synergy may reflect the emergence of task-specific, feedforward postural integration for throwing. Rather than a canonical anticipatory postural adjustment that precedes the focal action, our data suggest near-synchronous trunk–upper co-activation reflecting feedforward posture– action integration consistent with the developmental maturation of posture–action coupling (13–15).

Fractionation was also evident: a bilateral soleus-dominant synergy (PS-8) was divided into lateralized modules (SA-7 and SA-8), suggesting a functional specialization that may support asymmetric lower-limb contributions during throwing. This shift likely enhances force transfer along the kinetic chain and reflects growing specialization of lower-limb contribution to throwing (2). Additionally, PS-9, characterized by coactivation of the right rectus femoris and right soleus, may represent an alternative strategy contributing to force generation in participants adopting a two-footed stance.

Together, these findings demonstrate that merging and fractionation represent complementary mechanisms by which synergy organizations adapt to support developmental refinement of throwing performance.

### Functional role of each synergy

Despite these reorganizations, six synergies were commonly observed in both the preschool and school-aged groups. This persistence may support the notion, as proposed in earlier studies (2), that the ability to throw is an innate human trait, with fundamental primitives already established in early childhood. Below, we describe the possible functional roles of the observed synergies, excluding those described above, using conventional throwing phases as a reference framework rather than implying that children executed these phases in the same manner as skilled adult throwers. PS-1/SA-1, dominated by the right tibialis anterior, likely contributes to drive-leg ankle stabilization and balance at the initiation of the throwing motion (16, 17). PS-2/SA-2, characterized by co-activation of pectoralis major and rectus abdominis, may serve as a trunk– upper-limb “acceleration” module that aids force transmission toward ball release (18–20). PS-3/SA-3, involving the deltoid, infraspinatus, and trapezius, appears related to humeral elevation, scapulothoracic control, and eccentric stabilization during arm raising and deceleration (19, 21). PS-4/SA-4, comprising the right erector spinae and left biceps femoris, suggests lumbopelvic coupling that supports trunk rotation and posterior-chain involvement as the stride progresses (18, 22). Finally, PS-5/SA-5 (left tibialis anterior) and PS-6/SA-6 (left rectus femoris) are consistent with stride-leg stabilization and braking after landing, facilitating efficient energy transfer up the kinetic chain (22). Taken together, these modules appear to underlie essential functions such as drive-leg stabilization, scapular and humeral control, trunk–pelvis coupling, and stride-leg braking. Their persistence suggests that fundamental primitives for throwing are established by early childhood, providing a foundation upon which later refinements build.

### Refinement of temporal coordination

A key finding was the reduction in variability of synergy activation timing (i.e., CoA) in SA. This result contrasts with a previous study focusing solely on upper-limb muscle, which reported minimal age-related timing differences (7) and underscores the importance of analyzing whole-body coordination. In preschoolers, immature sequencing of trunk, limb and postural actions likely contributes to trial-to-trial variability (23, 10). For example, if trunk rotation begins too early or too late relative to stride-foot contact, or if arm acceleration is not well synchronized with trunk motion, the overall sequence becomes inefficient and unstable (24), leading to broader temporal dispersion of synergy activation.

Several factors may account for the developmental refinement of CoA. First, maturation and task-specific experience promote tighter synchronization of support actions, trunk rotation, and arm acceleration with key mechanical events such as stride-foot contact, peak trunk rotation, and ball release, thereby narrowing coordination solutions and reducing variability (23, 10). Second, the timing patterns of pelvic and trunk rotation, as well as lower-limb contribution, vary with both skill level and developmental stage, but with increasing maturity, these patterns converge toward a more stereotyped temporal sequence (25, 26). Consequently, school-aged children demonstrate more reproducible coordination, with synergy more precisely aligned to critical mechanical events (27, 28).

This reduction in timing variability reflects not only greater efficiency but also a protective mechanism: consistent sequencing minimizes abnormal joint loading, particularly at the shoulder and elbow (24, 29), and enhances smooth energy transfer along the kinetic chain (23, 30).

### Neural mechanisms underlying synergy reorganization

Muscle synergy organization can be flexibly reconfigured or fine-tuned to improve performance during development (5, 6), adaptation (31–33), and skill learning (34, 35). The reorganization observed here likely reflects converging influences of neural maturation, motor experience, and biomechanical growth. Neural factors such as myelination, increased synaptic efficiency, and refinement of corticospinal pathways, enable more selective control of muscle subgroups within existing synergies (36, 37), supporting both structural modifications and sharpening temporal activation patterns (6, 38). Experience through spontaneous play and repeated throwing likely promotes effective temporal sequences via trial-and-error learning and Hebbian-like plasticity, biasing synergy merging or fractionation toward task-effective compositions (39, 40). In parallel, biomechanical scaling during growth enhances stability and mechanical leverage, reducing the need for compensatory timing adjustments (41). Together, these processes drive selective synergy reorganization while preserving the core modular repertoire.

## Conclusions

Whole-body muscle synergies supporting overarm throwing are established early in childhood but undergo structural reorganization and temporal refinement between preschool and school age. These changes, particularly the integration of upper-limb and trunk synergies, fractionation of lower-limb synergies, and improved consistency of activating timing, likely underlie developmental improvements in throwing performance. Although the number of synergies remained unchanged in this developmental window, later stages may bring quantitative increases. Overall, early throwing skill enhancement depends less on expanding the number of synergies and more on refining their structure and timing, making an important stage in the broader trajectory of motor development.

## Materials and Methods

### Participants

11 preschoolers (PS; 5 males, 6 females; 62.2 ± 8.0 months, range 47–77 months) and 9 school-aged children (SA; 4 males, 5 females; 100.3 ± 14.4 months, range 85–131 months) participated in this study. None had current musculoskeletal injuries or a history of neurological impairment within the past year, or prior experience in sports involving overarm throwing (e.g., baseball or softball). The study was conducted in accordance with the Declaration of Helsinki, and all procedures were approved by the local committee in the University of Tokyo (reference number: 895-6). Written informed consent was obtained from the parents of all participants.

### Experimental tasks

All experiments were conducted in an indoor laboratory. Participants were instructed to throw a ball with maximum effort toward a net located 2 meters away. After a brief warm-up and familiarization, each participant performed 8 to 11 throws.

### Data recordings

Ball kinematics were recorded with two synchronized high-speed cameras (DSC-RX10M4, SONY, Japan; 960 fps) placed behind and to the side of the participant. Synchronization was achieved with an LED flash triggered by a commercial device (FA-WRC1M and FA-WRR1, SONY, Japan). The center coordinates of the ball were extracted from the recorded images without physical markers, using the same methods as in a previous study (42). Each recording window lasted ∼1000 ms, covering the throwing motion from initiation to several tens of milliseconds after ball release. Trials were excluded if the ball moved out of the camera’s view or if the LED light flash occurred outside the data acquisition window due to motion variability.

We simultaneously recorded EMG activity from 16 muscles spanning the upper limbs, trunk, and lower limbs using wireless sensors (Pico EMG, Cometa Srl, Italy). Recorded muscles included pectoralis major (PM), middle deltoid (MD), infraspinatus (ISP), and middle trapezius (MT) on the throwing side; rectus abdominis (RA), erector spinae at the T10 vertebral level (ES/T10), rectus femoris (RF), biceps femoris (BF), tibialis anterior (TA), and soleus (SOL) bilaterally. EMG signals were sampled at 2000 Hz. Synchronization with video data was achieved using the LED flash signal.

### Kinematic analysis

Ball tracking in the high-speed camera images was performed using DeepLabCut (43), a deep learning-based automatic image recognition system. The tracking models were trained using manually labeled images from one trial per participant. For each trial, 20 frames were automatically selected by built-in frame extraction algorithm of DeepLabCut, which uses the k-means method to identify the most informative frames for manual labeling. Separate models were developed for each camera view and trained for 100,000 iterations. These trained models were used to automatically track and digitize the center of the ball in all remaining trials. This tracking procedure was identical to that used in previous studies involving throwing tasks (42, 44). The accuracy of this method has been validated against conventional methods, with the absolute distances in coordinates average ranging from 15.5 to 29.4 mm, and correlation coefficients between the two methods ranging from 0.932 to 0.999 (44).

Two points during the throwing movement, Start and Ball release, were detected using the position coordinates of the ball, and the analysis window was defined as the phase from Start to Ball release. Start was defined as the frame where the ball speed exceeded 5 km/h, while Ball release was determined by qualitative observation of the images from the side-view camera. Ball speed was calculated as the magnitude of the velocity vector at the moment of ball release.

### EMG processing

EMG data were processed using custom MATLAB script (R2024a; MathWorks, Natick, MA, USA). Signals were band-pass filtered 40 and 450 Hz, demeaned, full-wave rectified, and low-pass filtered at 10 Hz (Finite Impulse Response filter, or FIR) (6, 45). Subsequently, each muscle’s amplitude was normalized to its peak value within each trial (46, 47). Data were then time-interpolated to 201 points per trial, producing the EMG matrix of 16 muscles × 201 data points. For each participant, trials were concatenated in the time dimension, resulting in a two-dimensional EMG matrix of 16 muscles × (201 data points *× n* trials) for muscle synergy analysis (48, 49).

### Muscle synergy extraction

Muscle synergies were extracted using non-negative matrix factorization (NMF) (50, 51) decomposing EMG into muscle weighting (W) and temporal patterns (C):

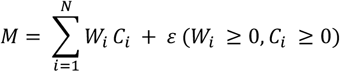

where *M* is the EMG matrix (muscles × time points), *W* denotes muscle weighting representing the contribution of each muscle to each synergy, and *C* represents a temporal pattern indicating the time-varying activation coefficients of each synergy. Both *W* and *C* contain only non-negative values, reflecting the physiological constraint that muscle activations cannot be negative. This decomposition allows the identification of a small number of underlying muscle synergies that can reconstruct the original EMG patterns with minimal error.

The number of synergies was determined based on the Variance Accounted For (VAF), which quantifies how well the reconstructed EMG signals (*W*×*C*) approximate the original signals (*M*). VAF was calculated using the following formula:

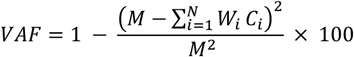

where the summation is taken over all elements of the matrix. We selected the minimum number of synergies such that the VAF exceeded 90% (49), ensuring that the decomposition retained the essential features of the original EMG signals while reducing dimensionality.

### Clustering muscle synergies

To characterize differences in synergy structure between the two groups, representative synergy vectors were first identified within each group using *k*-means clustering. Clustering was performed in MATLAB using the k-means function, with the number of clusters ranging from 2 to 12 (6). The algorithm was set to run with a maximum of 500 iterations per clustering attempt and repeated 3000 times with different random initial centroids to ensure robustness and avoid local minima. The optimal number of clusters was determined based on the average silhouette score (33, 52), and the centroid of each cluster was interpreted as a representative synergy pattern within the group. Differences in the distribution of synergy types between the PS and SA groups were then examined to explore developmental changes in muscle coordination.

### Muscle synergy similarity

To quantitatively assess the similarity of muscle synergies between the PS and SA groups, we calculated pairwise cosine similarity between synergy vectors (53). Each synergy vector was normalized to unit length prior to comparison to ensure that only the direction (i.e., muscle weighting pattern) was considered independent of magnitude. Cosine similarity was computed using the following formula:

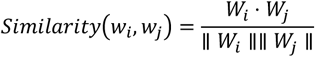

where *W*_*i*_and *W*_*j*_ represent two synergy vectors and denotes the dot product. A similarity value of 1 indicates perfect similarity, whereas values closer to 0 indicate dissimilarity. To identify shared muscle synergy patterns between developmental stages, we compared synergy vectors across groups (PS vs. SA). Synergy pairs with a cosine similarity ≥ 0.9 were considered to reflect similar muscle coordination structures (53). The proportion of highly similar synergy pairs was then calculated to evaluate the extent of similarity between the two groups.

### Merging of muscle synergies

To evaluate developmental changes in synergy structure, we analyzed the merging of muscle synergies between groups (54). In this study, merging was defined as the case in which a synergy vector observed in SA group could be reconstructed as a linear combination of two or more synergy vectors from PS group. Formally, this relationship is expressed as:

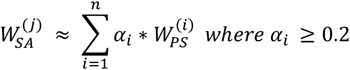

Here, 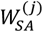 denotes a synergy vector from the SA group, and 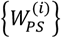 are synergy vectors from the PS group. The coefficients *α*_*i*_ were estimated using non-negative least squares. To identify meaningful merging, we applied an additional threshold: only synergy vectors with *α*_*i*_ ≥ 0.2 were considered as contributing to the merged synergy. If two or more PS synergies met this criterion and the reconstructed vector showed high similarity to the original SA synergy (cosine similarity ≥ 0.9), the case was classified as a merging event. The number and proportion of such merged synergies were calculated to quantify the extent of integration. This merging process was interpreted as a developmental consolidation of motor modules, potentially reflecting increased efficiency and flexibility in neuromuscular control.

### Fractionation of muscle synergies

To evaluate how muscle synergies change during early development, we analyzed whether individual synergies observed in PS group could give rise to multiple distinct synergies in SA group. This process, referred to as *fractionation*, was defined as the case in which a synergy from the PS group could be approximated by a linear combination of two or more synergy vectors from the SA group (54). Formally, this relationship is expressed as:

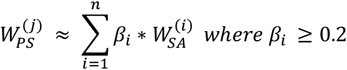

Here, 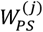 denotes a synergy vector from the PS group, and 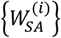 are synergy vectors from the SA group. The coefficients *β*_*i*_ estimated using non-negative least squares. To identify meaningful cases of fractionation, we considered only those in which two or more SA synergies contributed substantially to the reconstruction (*β*_*i*_ ≥ 0.2), and where the reconstructed vector showed high similarity to the original PS synergy (cosine similarity ≥ 0.9). The number and proportion of such fractionated synergies were then calculated as an indicator of developmental differentiation of motor modules. This fractionation was interpreted as reflecting increased flexibility in neuromuscular control, enabling finer motor adaptation as the neuro-musculoskeletal system matures.

### Temporal analysis of synergy activation

For each synergy identified in each participant, the centroid of activity (CoA) defined as the center of gravity of the corresponding temporal pattern was calculated using the following equation:

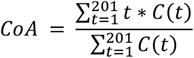

where *C*(*t*) represents the temporal pattern at normalized time frame *t* (201 frames per trial). To facilitate interpretation, CoA values were linearly rescaled from frame indices to a percentage scale (0–100%), such that 0% corresponds to Start and 100% to Ball release time.

This metric was employed to objectively characterize the temporal distribution of each synergy (55). By weighing each time point according to its activation magnitude, CoA reflects the overall timing and structure of activation throughout the movement phase, rather than focusing on a single peak value. Furthermore, by calculating the standard deviation of CoA across repetitions, we were able to assess the consistency of activation timing, which serves as an index of movement stability. We quantified timing stability at the participant level as SD of CoA, defined as the mean of the per-synergy standard deviations of CoA calculated across repetitions within that participant.

### Statical analysis

All statistical analyses were performed using MATLAB (R2024a; MathWorks, Natick, MA, USA). Independent samples t-tests were conducted to compare ball speed, the number of extracted muscle synergies, and the standard deviation of the CoA between groups. Data normality was assessed using the Shapiro–Wilk test, and homogeneity of variances was evaluated using Levene’s test. Statistical significance was set at *p* < .05.

## Acknowledgements

We thank the Doronko Welfare Foundation for their support in participant recruitment and for facilitating the smooth execution of the experiments. We are also grateful to Dr. Ken Takiyama (Tokyo University of Agriculture and Technology) for kindly providing the equipment used in this study.

## Funding

This work was supported by the Nippon Professional Baseball (NPB) Organization and AGEKKE CORPORATION.

## Notes

### Competing Interest Statement

The authors have declared no competing interest.

